# Parameter estimation in mathematical models of viral infections using R

**DOI:** 10.1101/130674

**Authors:** Van Kinh Nguyen, Esteban A. Hernandez-Vargas

## Abstract

In recent years, mathematical modeling approaches have played a central role to understand and to quantify mechanisms in different viral infectious diseases. In this approach, biological-based hypotheses are expressed via mathematical relations and then tested based on empirical data. The simulation results can be used to either identify underlying mechanisms, provide predictions on infection outcomes, or evaluate the efficacy of a treatment.

Conducting parameter estimation for mathematical models is not an easy task. Here we detail an approach to conduct parameter estimation and to evaluate the results using the free software R. The method is applicable to influenza virus dynamics at different complexity levels, widening experimentalists capabilities in understanding their data. The parameter estimation approach presented here can be also applied to other viral infections or biological applications.

## 1. Introduction

Seasonal epidemics and pandemics of influenza virus infections remain a major health burden worldwide, causing immense losses in lives, life quality, and economy (1-3). The overwhelming amount of influenza research has largely improved our understanding, however, holistic understanding that promotes serious adverse events leading to health complications are largely fragmented (4).

Analyses of experimental data on viral infections have predominantly based on statistical methods. These approaches assist experimentalists to recognize differences and correlations, but in-depth interpretations of the underlying mechanisms are limited. With mathematical modeling approaches, one can formulate different hypothesized mechanisms in forms of mathematical relations. Consequently, parameter estimation procedures are performed to test the models against empirical data (4-7). This method has been used to study a wide range of events occur during the progression of influenza infection (4-6,8). For instance, mathematical models have been used to describe the viral replication cycle, interactions between the virus and the host, and the outcomes of the infection (Figure 1). Additionally, simulation results can reveal not only the basic characteristics of the infection dynamics but also practical knowledge in controlling the infection (4-6,9).

**Figure 1.**
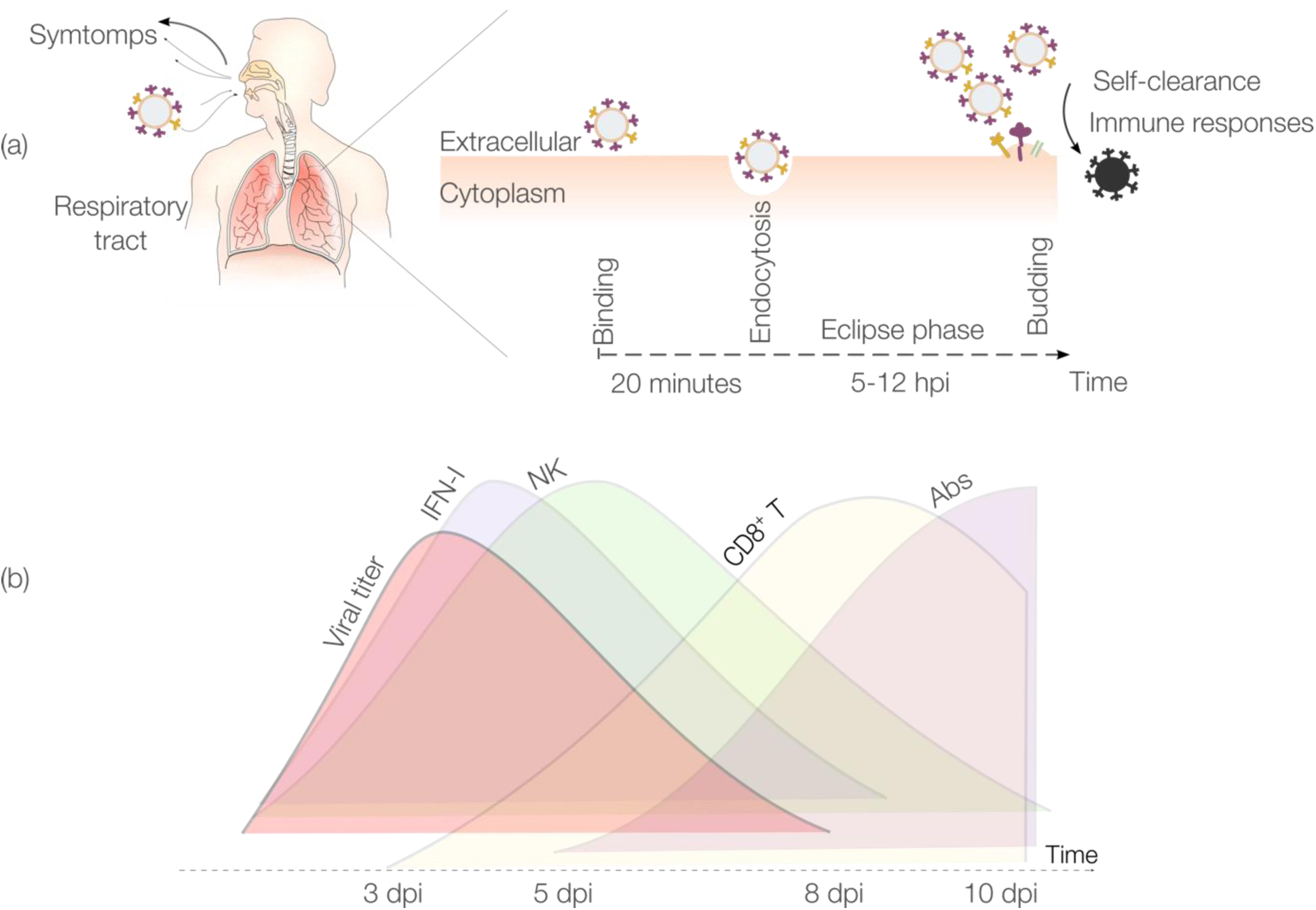
IAV infection and dynamics. (a) Description of the main phases of IAV infection within a host. After entering the respiratory tract, each virion binds to a target cell. Then, virions enter the eclipse phase (5-12 hpi), before starting to replicate and infecting other cells. (b) IAV and IR dynamics. The innate IR is mainly represented by Type I interferon (IFN-I) and by the natural killer (NK) cells, whereas the adaptive IR is mainly driven by cytotoxic CD8+T cells (CTLs) and antibodies (Abs). Figure adapted from (4).

Conducting parameter estimation for mathematical models, however, is a demanding task. This requires familiarizing with different concepts of mathematics, optimization, programming language, and sometimes costly software toolboxes. Nevertheless, these technical problems should not prevent biologists and virologists from exploring their data potentials. Thus, in this chapter, we introduce an adaptable and state-of-the-art protocol for parameter estimation and evaluation. To this end, we focused on ordinary differential equations (ODEs) to model the infection dynamics. The target-cell limited model presented in (10,11) is adopted owing to its role as the core component of more than a hundred publications in virus research, e.g., influenza (4,5,7,9,12), HIV (13,14), and Ebola (15) among others.

This chapter covers chronologically the steps portrayed in Figure 2. Briefly, experimental data need to be prepared in standard formats. Model equations need to be defined with relevant components and corresponding model parameters. Based on that, a cost function that defines how matching the model and the data is written in the R programming language (16). To this end, the root mean square errors function is considered. The function, the data, and the model can be then fed into an optimizer algorithm to find the best set of parameters that provide the best agreement between the model and the data. The global optimization named Differential Evolution (17) is used here to adjust the model parameters. Setting conditions of the optimizer and the plausible range of the parameters of interests need to be defined. When contradictory hypotheses exist, model comparison among them can be done at this point with information criteria. Then, model predictions can be performed with the obtained parameters. Further model evaluation steps are developed using the best model to obtain confidence intervals or to detect potential drawbacks of the obtained parameters.

**Figure 2.**
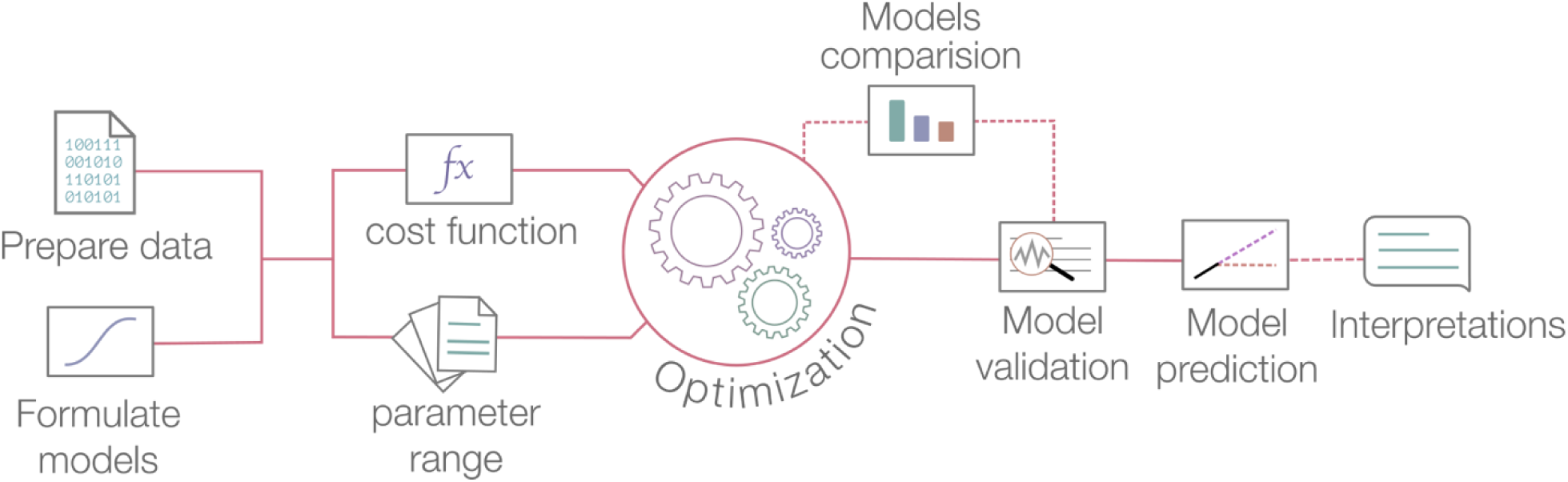
A typical parameter estimation process in mathematical modeling. Dashed lines indicate optional steps and those are not presented in the scope of this chapter.

## 2. Materials

The following materials are needed:

1. Experimental data: For illustration purpose, we consider a synthetic *in vitro* data set of influenza A virus infection with the viral dynamics and the sampling scheme resemble that in (18). Approximately 106 host cells are assumed to support influenza virus infection. Based on practical lab limitations, the viral load is assumed to be the only measurement, thus, viral titers were measured regularly (in TCID_50_) at day 1, 2, 3, 5, 7, 9 post infection. At each time point, there are five replications. The data used in this chapter can be found in this external hyperlink. The first term in each row represents the sampling date and the second is the viral load. Note that viral titers were already converted into log base 10 scales (see Notes 1).
2. Mechanistic model(s): The proposed mathematical model depends on the data at hand and the hypothesis to be addressed. Here, we are interested in having the estimates of the rates of infection, infected cell death, viral replication, and viral clearance. Thus, a widely used model for viral infection so-called the target-cell limited model (10) can be used (Figure 3).

**Figure 3.**
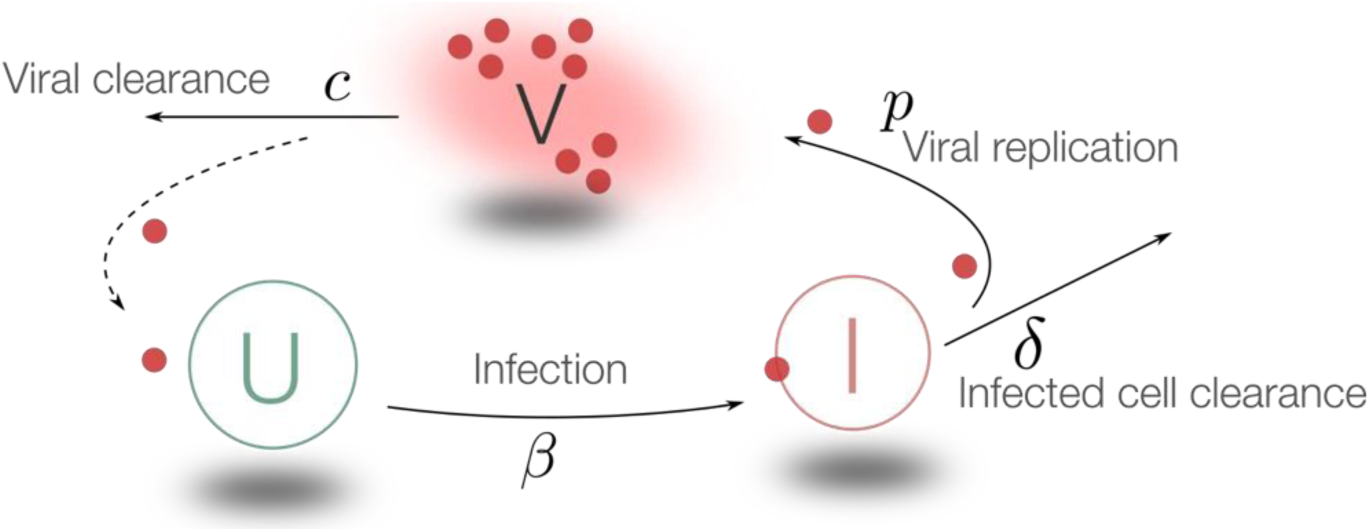
Schematic of the target-cell limited model. This model assumes that viral infection is limited only by the availability of the uninfected cells. The roles of the immune systems are neglected. The uninfected cells (U) are infected by the viruses and become infectious (I), consequently, these infectious cells are able to release virus particles (V). The viruses can continue to infect the remain susceptible cells.

 The model includes three compartments: uninfected cells (U), infected cells (I), and viral titers (V). The model reads in the following three differential equations:

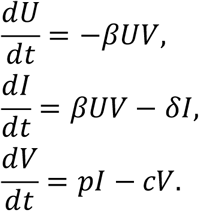
The left term of the equations represents the change of the variables respect to the time. The parameters *β*, *δ*, *p*, and *c* represent the rates of effective infection, infected cell death, viral replication, and viral clearance, respectively. It is considered that the virus (V) infects susceptible cells (U) with a rate *β*. Infected cells are cleared with a rate *δ.* Once cells are productively infected (I), they can release virus at rate *p* and virus particles are cleared at rate *c*.
3. A computer with any of the following operating systems: Windows, Linux, or Mac OS.
4. R software: a free, open-source, and high-level programming language (16), downloading from https://www.r-project.org.
5. Required R’s packages include *deSolve* (19) (solving differential equations) and *DEoptim* (20) (performing the Differential Evolution algorithm) can be installed in R by running the following commands§

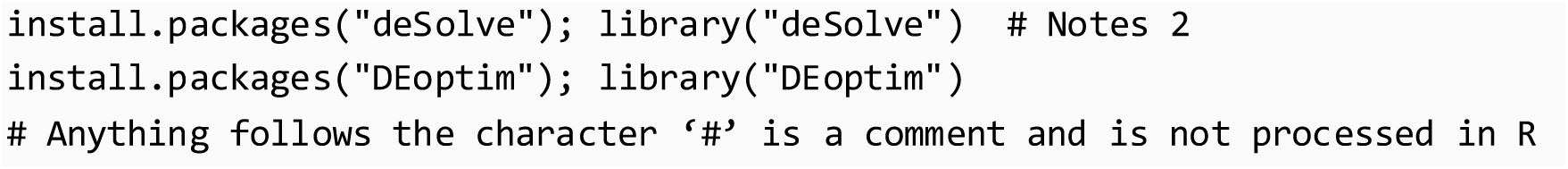

For the rest of the chapter, fix-width font-style letters denote R codes.

## 3. Methods

### 3.1. Preparing data

The experimental data stored in Excel sheet are most often not ready for analyses in R. Comma-Separated-Values (.csv) is a universal format that can be read in any software. This can be achieved by:

1. Deleting all irrelevant data (notes, comments, etc.) in the Excel sheet,
2. Naming variables (columns) with computer-friendly format, i.e., no spaces or special characters, starting only with characters not with numbers,
3. Choosing *Save As*, in file format field choose *Comma Separated* (see Notes 3, 4).

Then, reading the data into R can be done by running:

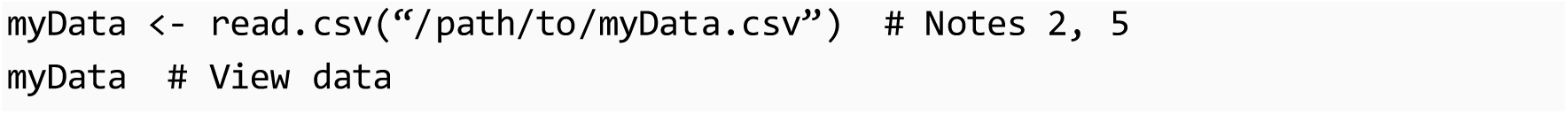

The data have two variables (each in different column), including *V* (viral titters in log base 10) and the time (in days). For simplicity, we can also input directly the data as follow:

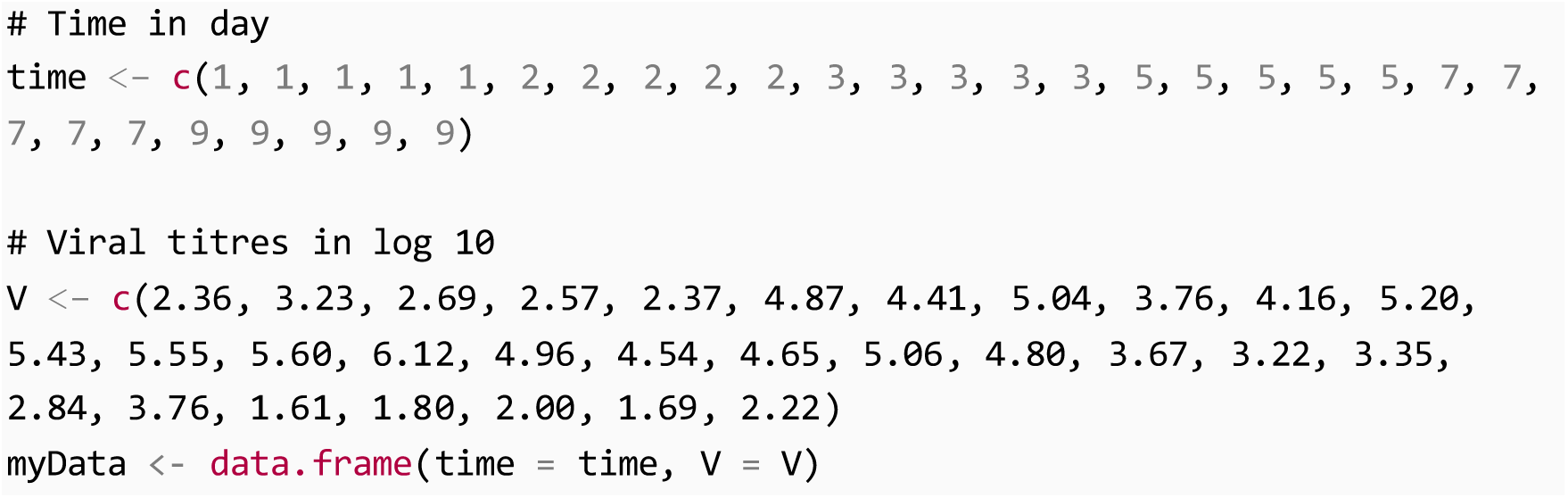

### 3.2. Writing the model and the cost function

1. The target-cell limited model can be written in R format as follows:

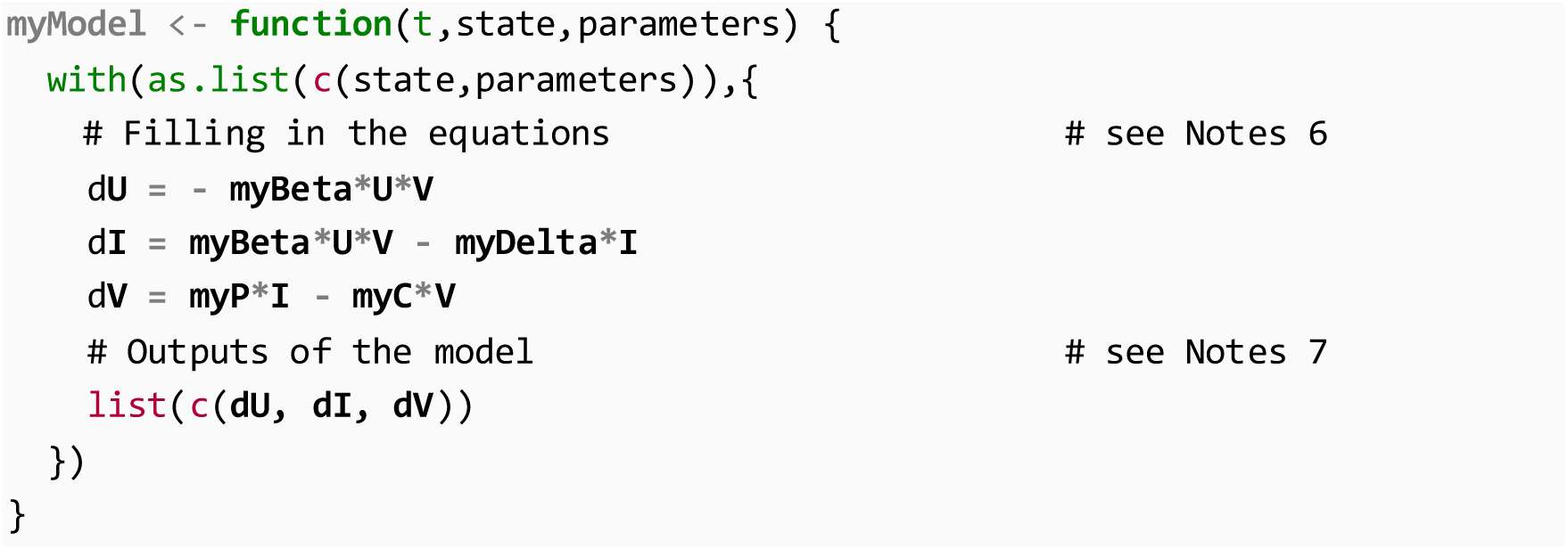

In short, the model is named as *myModel*. The three components *U*, *I* and *V*, called states in modelling terms, are written in separate lines with leading letter *d*. The right-hand side of the equations are the same as the model equations, except the variable names are spelled out (see Notes 6). We also define what is important in the model for later uses, including and the time scale for the model. This can be arbitrarily chosen but it should cover the range of the observed time scale. Here we record the model outputs approximately every 15 minutes during ten days.

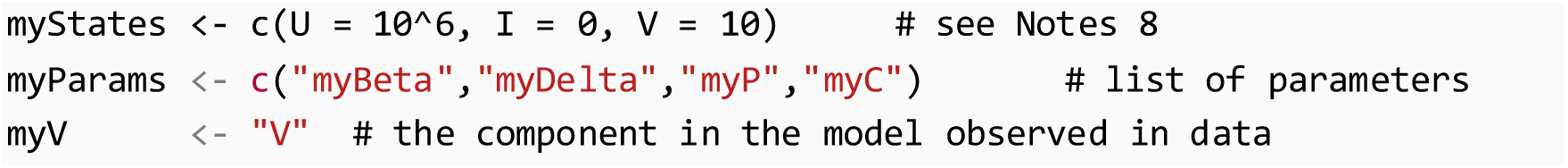

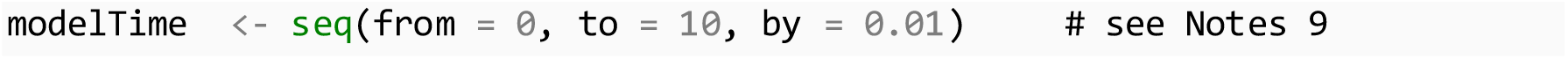
2. The root mean square errors (*RMSE*) measures the magnitude of the difference between the output from the model (*V*) and the experimental data. Here it is calculated as

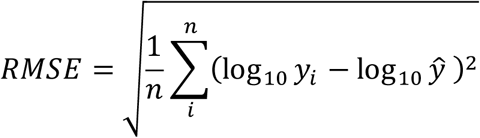

where *n* is the sample size, *y*_*i*_ is a data point, and ŷ is the value produced from the model when we plug in a set of parameter values and calculate the value of *V* at the time point observing *y*_*i*_. In R language, the cost function can be written as

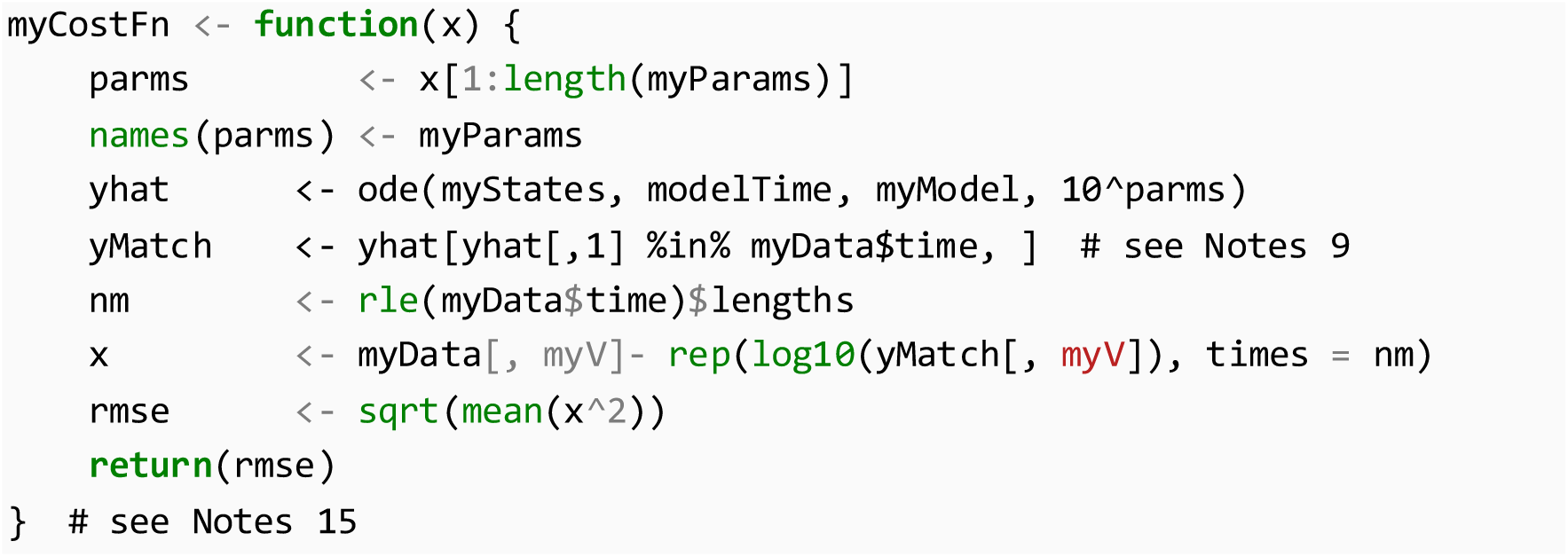

#### 3.2.1 Defining parameter boundaries, optimizer conditions and running optimizer

1. Based on literature, we define ranges of plausible parameter values. For example, the elimination half-life (*t*_1/2_) of influenza is unknown, however this cannot be considered either in seconds or in a century time scale. Converting this time into the clearance rate *c* is done by the formula *t*_1/2_ = ln(2) /*c*. Note that the boundaries are transformed to log base 10 scale and that the numbers subject to one’s own expertise. We let the reader check the boundaries of the four parameter *β*, *δ*, *p*, and *c* below.

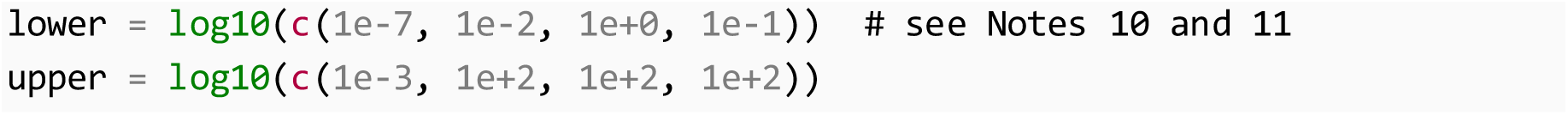
2. Standard optimizer settings are often not sufficient for complex models. We can force the optimizer to work more exhaustively by increasing the number of trials (combination of *itermax* and *steptol*) and decreasing the relative tolerance named as *reltol* (measurement of the error relative to the size of each solution component).

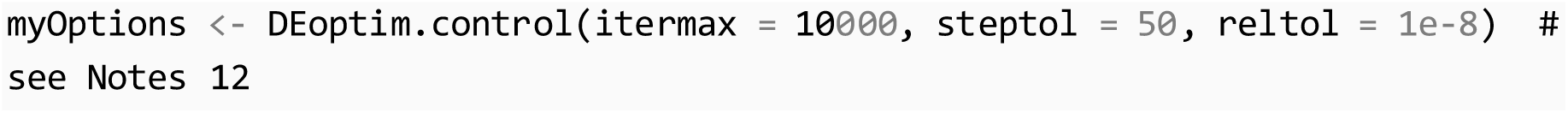
3. At this point, we are ready to fit the model by calling the optimizer with the inputs including the cost function, the lower and upper bound, and the options as follows:

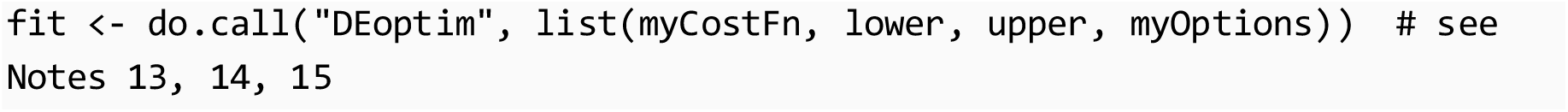

The simulation time took approximately five minutes on an Intel Core i7, 8Gb RAM computer. The algorithm evaluates different combinations of the parameters in the provided ranges, comparing them by the *RMSE*. The process is repeated until the algorithm can not find a combination of parameters that has a significant improvement compared to the current parameter set.
4. Visualization of the results can be done with the following commands (Figure 4)

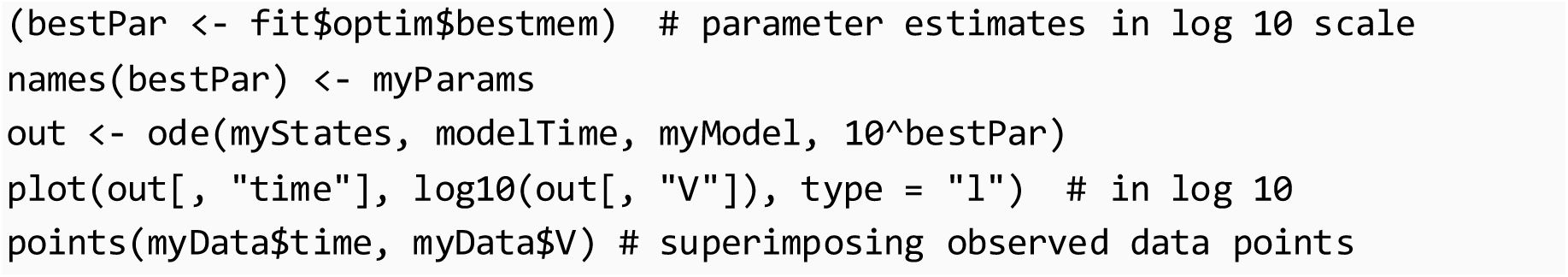

**Figure 4.**
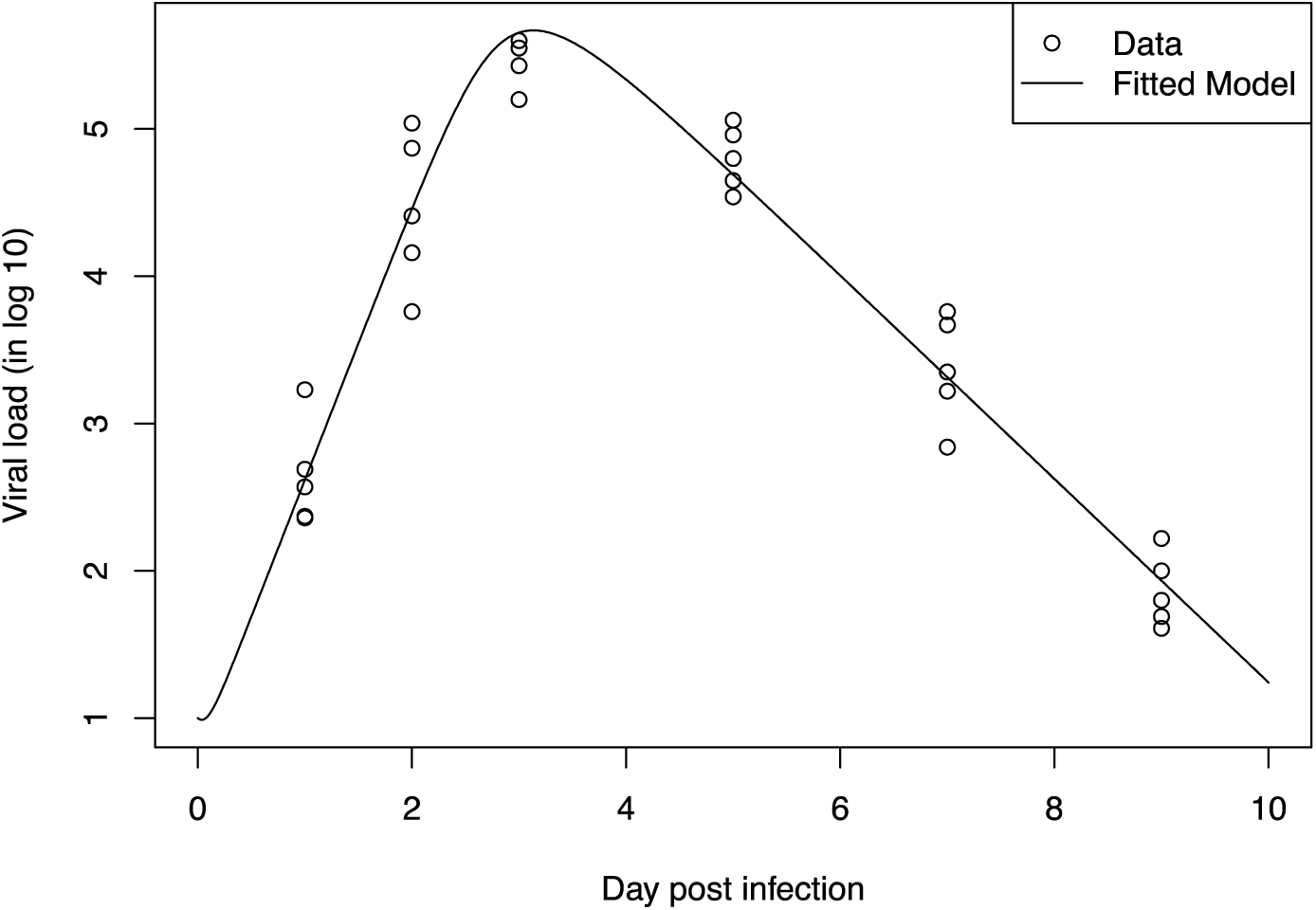
Fitted model and the observed data.

### 3.3 Model comparison with Akaike information criteria (AIC)

AIC is a criterion of choice when coming to comparing models (21,22). More complex models (more parameters) tend to provide a better fit, thus the AIC gives a penalty to the number of parameters to avoid overfitting (23). The smaller the AIC the better the model. In this context, a function to calculate can be defined as:

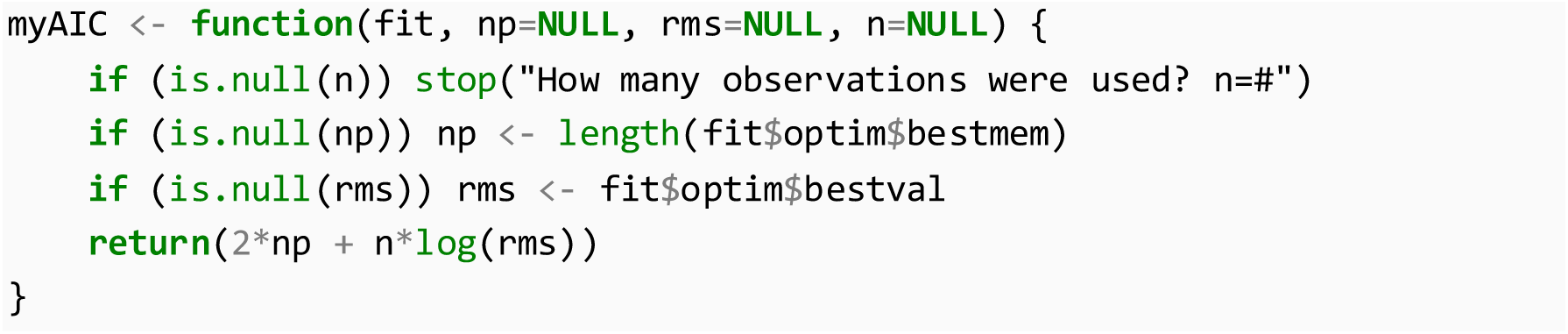

Computing the AIC by simply run:

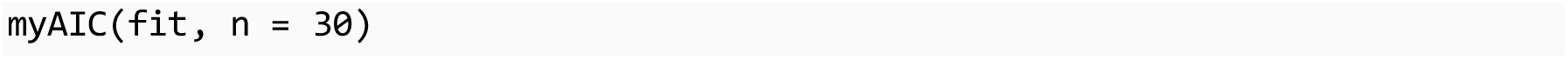

where *fit* is the fitted model and *n* is the sample size used for fitting (see Notes 16).

### 3.4 Likelihood profile of the model parameters

Because of the complexity of the model and the scarce of experimental data, it might be impossible to identify the parameters with less uncertainty (24,25). The profiling of model parameters is used to inform the identifiability of parameters or to calculate the parameter’s confidence interval. For each parameter, a sequence extending both sides of the estimated parameter to the boundaries is generated. For each value, the optimization is done by keeping the parameter value fixed while reoptimizing the other parameters.

1. Define the profiling function as below:

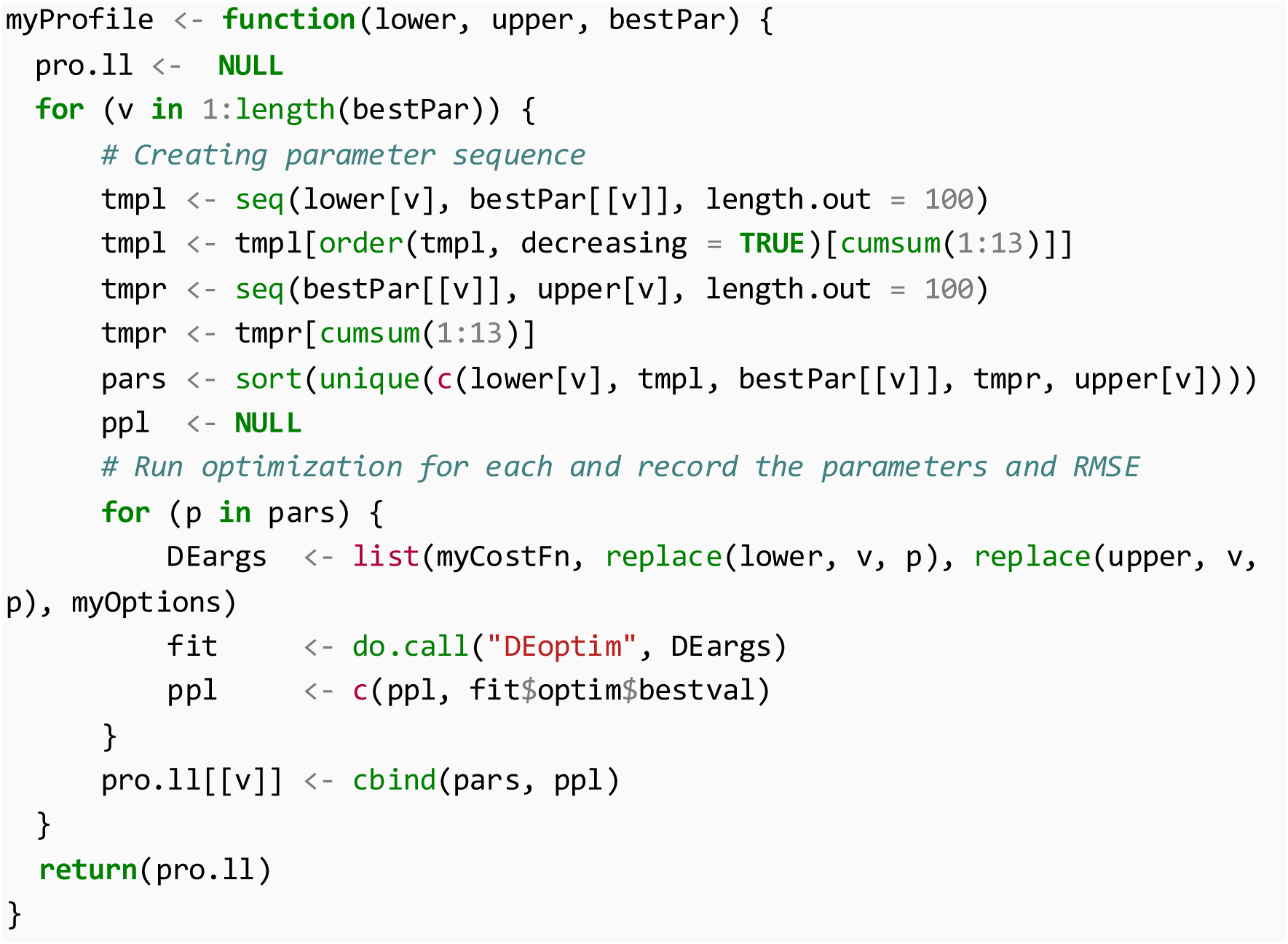

For efficiency, the parameter sequences were chosen such that it is dense close to the estimated values and is sparse towards the boundaries.
2. The profiling can now be executed simply by executing the following

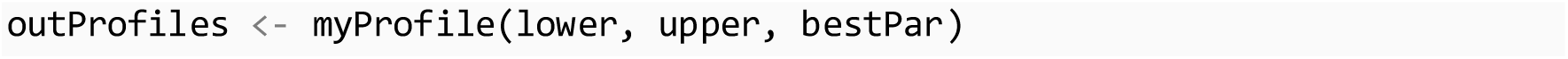

For each of the parameters, the previous function calculates 26 discrete points to form the likelihood profile. In total, more than a hundred optimizations will be run which could take hours or even days depending on the power of the computer (see Note 17).
3. Plotting the profile of the first parameters can be done by executing

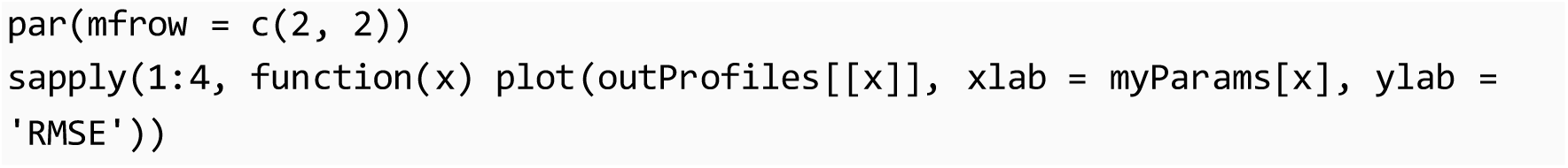

### 3.5. Bootstrapping parameters

Bootstrapping is a statistical method for assigning measures of accuracy, such as confident interval, to the parameter estimates (26-28). Here a nonparametric approach using Monte Carlo resampling is employed. First, we resample the data with replacement to have a sample of equal size to the original data. The parameters are then estimated from the resampling. The procedure is repeated many times to get bootstrap distributions of the parameters.

1. Define bootstrapping function as below

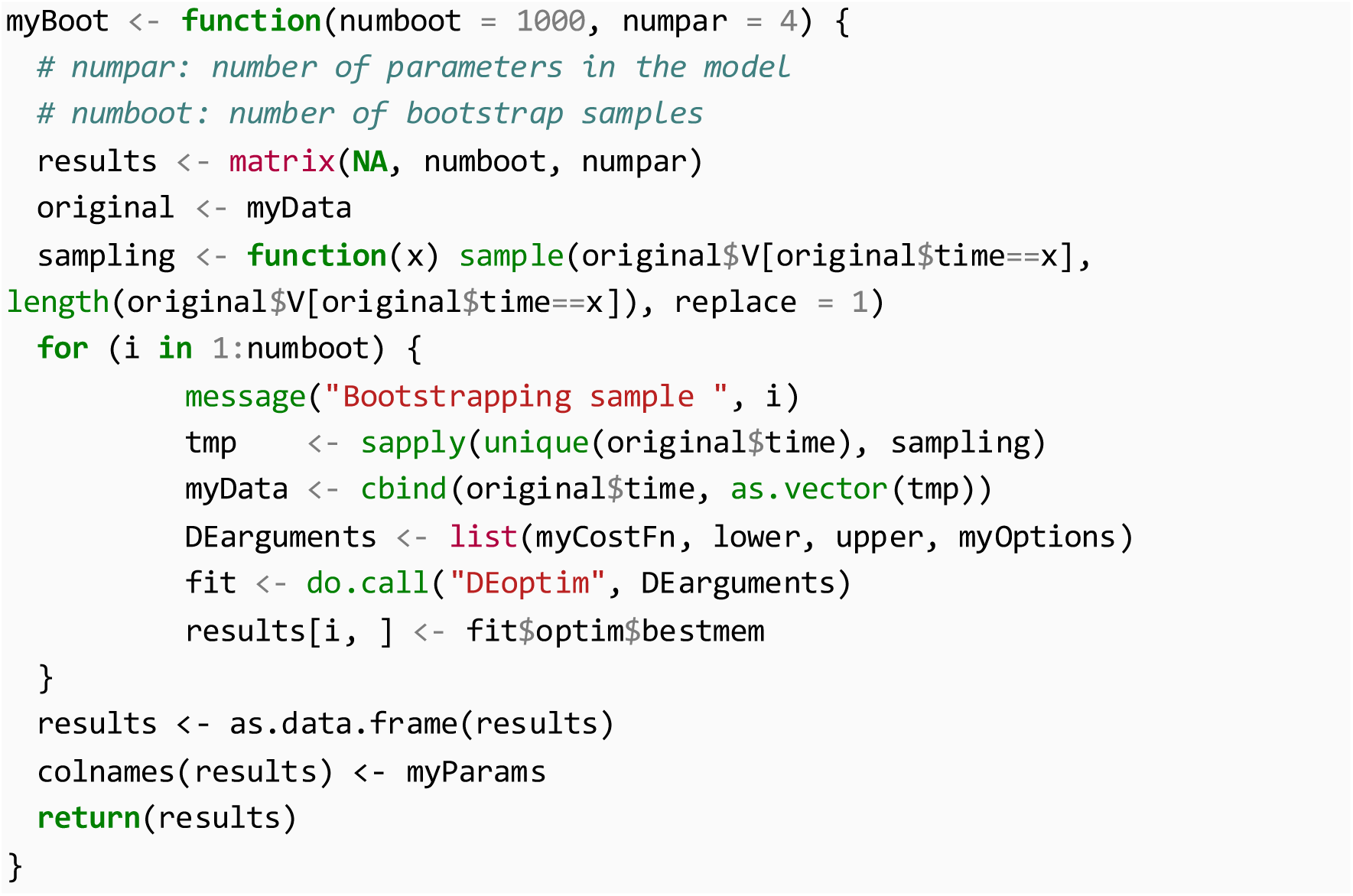
2. Running bootstrapping now can be done by executing

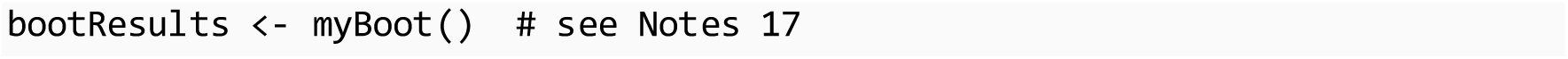

If you need to change the number of bootstrap samples, simply input the parameter *numboot*, e.g., *myBoot(10000).* Plotting histogram of the bootstrap samples can be done with the commands:

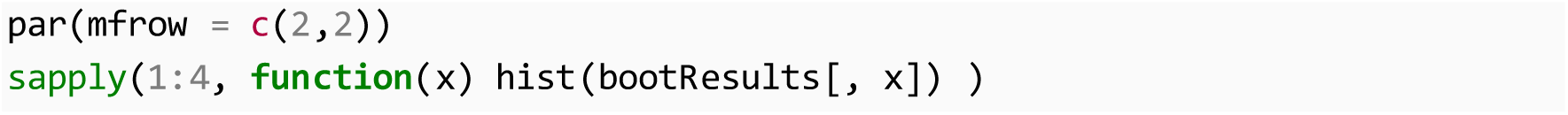
3. 95% confidence intervals can be calculated from bootstrap sample with the percentile method as:

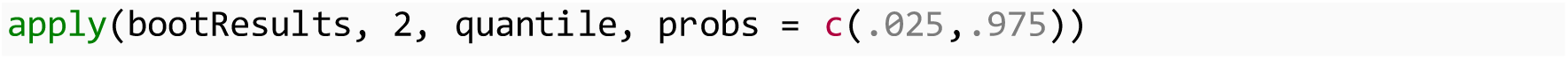

The functions and R codes presented in this section will work for other models and data sets as long as the same procedure and naming convention have been carried out.

## 4. Notes

1. The log-scale viral titers were used not only because of its conventional usage in reporting viral load, but the log-scale also assumes implicitly that viral load is normally distributed in log scale. This assumption simplifies the maximum likelihood problem to least squares (26), and thus the use of the *RMSE* as a cost function in the optimization.
2. R is a case-sensitive language. Check spelling carefully and whether the letters are properly capitalized.
3. To avoid unnecessary errors, CSV data files should be filled from the first row and first column, i.e., the first row for variable names and first column contains data of the first variable. R can read a wide range of data types and even directly from Excel, but specific functions are needed.
4. Double-checking data with decimal separators. Differences in locales of the operating system lead to wrong interpretation of number when saving as CSV. For example, a number stored in Excel as 3,141.2 would be potentially treated as two numbers: 3 and 141.2. This can be detected by visualizing the data in R or inspecting the CSV file directly to see how the data are interpreted. To avoid, numeric cells in Excel should be format without “*Use 1000 separator”* option enabled.
5. On the operating system of Windows, the path to the data file needs a special treatment in R, e.g., a file located in

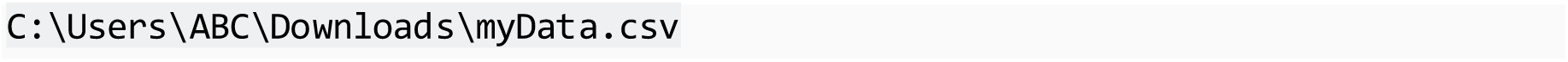

needs to be supplied to R as

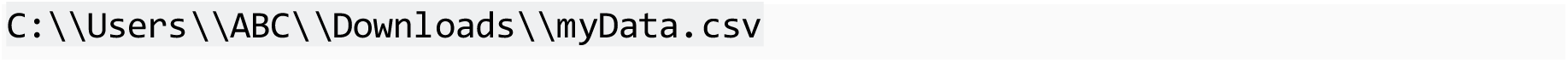

This is because a backslash is also an escape character in R.
6. If you see an error that says

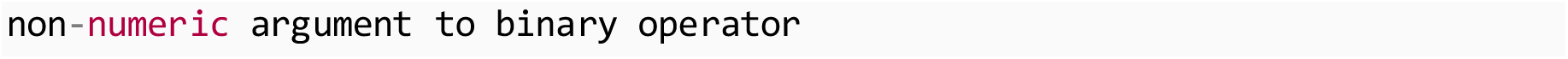

it is most likely that the name you gave to a model parameter already existed in R’s default environments, e.g., an expression as *–Beta* could lead to an error while *–myBeta* would not.
7. As noted by the deSolve package’s author (19), the order of the return values in *deSolve* model function is important. They need to strictly follow the order listed by the model equations written above it.
8. The model initial conditions are the values of each variable in the model at day zero. Although we optimize the model parameters in log scale, the model works in normal scale, and hence the initial conditions. Here, the initial number of the target cells was approximated at 106 from based on the experiment reports, the initial number of infected cells was zero. The initial viral titers were set at 10 TCID_50_. This value was arbitrarily chosen at a value below the detection level of 50 TCID_50_. To our knowledge, there is not any conclusive method to define this number while its value can affect the parameter accuracy (24). Smoothing and extrapolating approaches have been used (15) and seem to provide a reasonable estimate of the initial viral titers (24).
9. Because the model runs in a smoother time scale than the observed data, we calculated only the differences among the matched data points by the observed time. However, solving the ODEs are not always easy when we are evaluating large combinations of the parameters, some of which might not make sense at all. Thus, there might be cases when the exact time points cannot be computed but only its nearby neighbor. Matching these time points could lead to unexpected results. It is safer to truncate the numeric time points when matching the model time points and the data time points by replacing

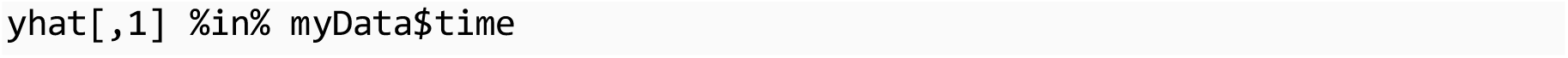

with

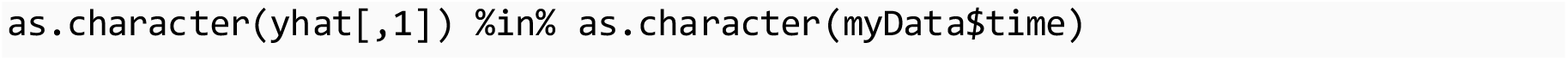
10. Some model parameters have analytical meanings but no equivalent experimental values. For example, the parameter *β* represents the reaction rate between the virus and the target cells, which depends on the concentration of the virus and the number of target cells. In these cases, the parameter’s range was defined to cover several orders of magnitude. Further insights into the choice of parameter ranges can be revealed by computing the parameter’s likelihood profile in section 3.4 (15,24,29).
11. Some model parameters can be determined experimentally. For example, the viral clearance rate *c* could be approximated by monitoring only the virus *in vitro* (30). In this case, we can reduce the burden on the optimizer and minimize the changes to the code by providing the same upper and lower bound for the parameter *c* at the formerly determined values, e.g.,

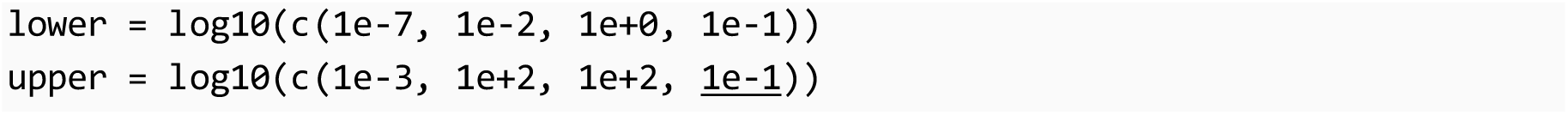
12. The differential evolution algorithm can be speed up considerably with parallel mode enable with the option such as (17):

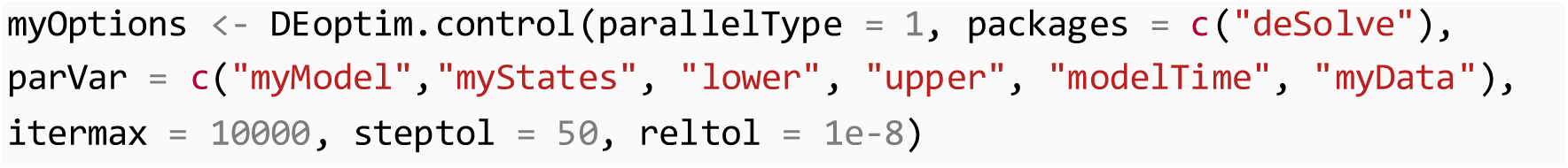

However, successful application of this for all calculations are not guarantee. Certain understandings about parallelization computing in R are needed to avoid miscalculations, e.g., wrong data or variables are used in calculation.
13. If we observed no changes in the cost function output (*bestval*) after several iterations, or even after reaching the maximum number of iterations (*itermax*), it means that the optimization failed. We can try to adjust the parameter boundaries to a more probable region based on literature and the parameter’s meaning.
14. As noted by the author (19), if you see an error as

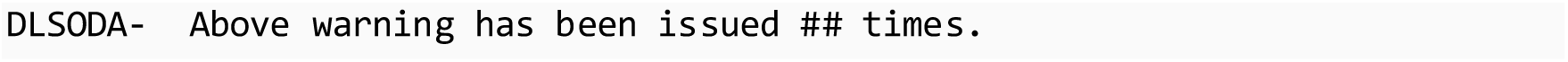

it means that the ODEs solver could not proceed with the current set of parameters. The most likely reason is that the parameter range was too wide, leading to extreme values to be evaluated. It can be avoided by narrowing down the parameter ranges to a more plausible region.
15. The implementation of the cost function in this chapter was a simplified version. More sophisticated error handling codes can be added to show what kind of error took place and how to handle it. This helps to speed up the process as well as prevents unexpected results that we are unaware of. This is done by the typical try-catch syntax, e.g.,

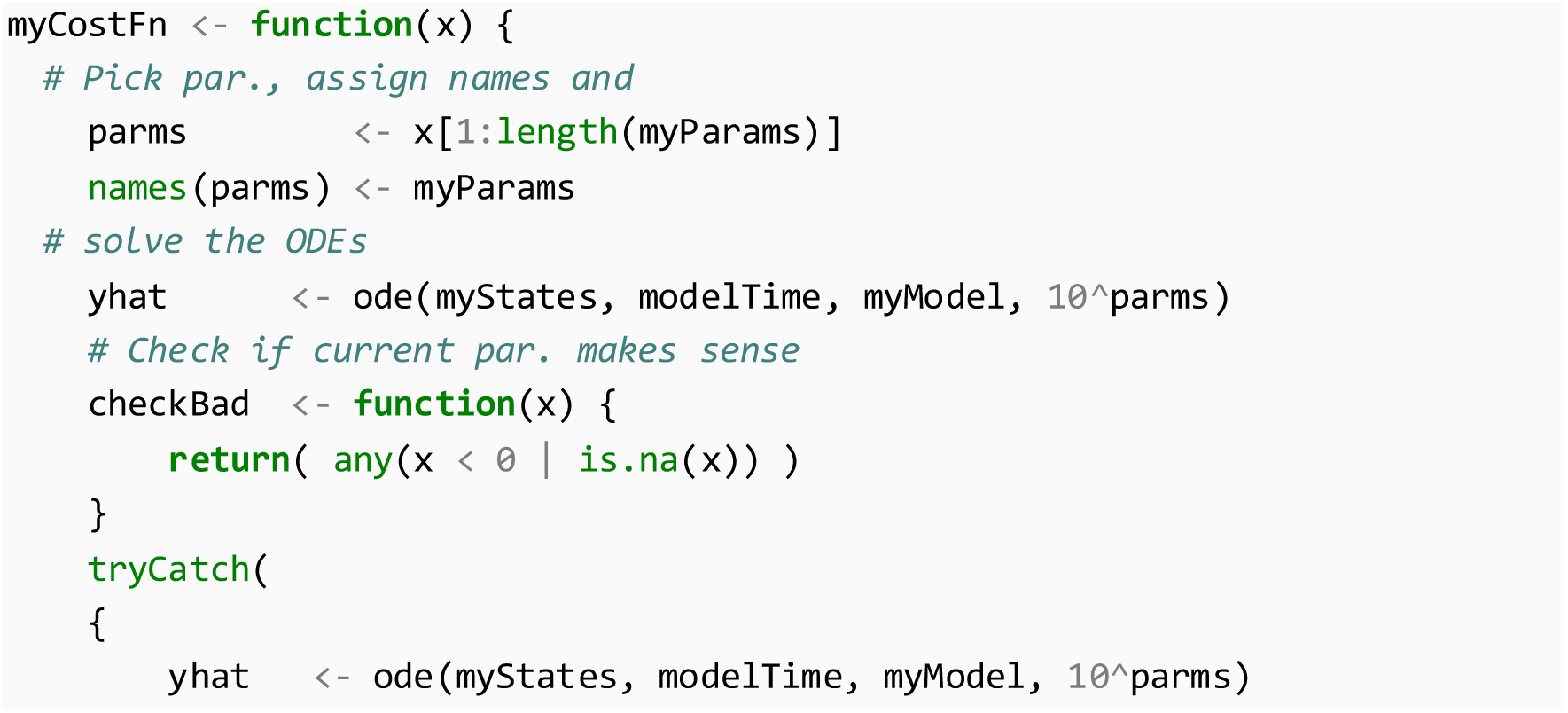

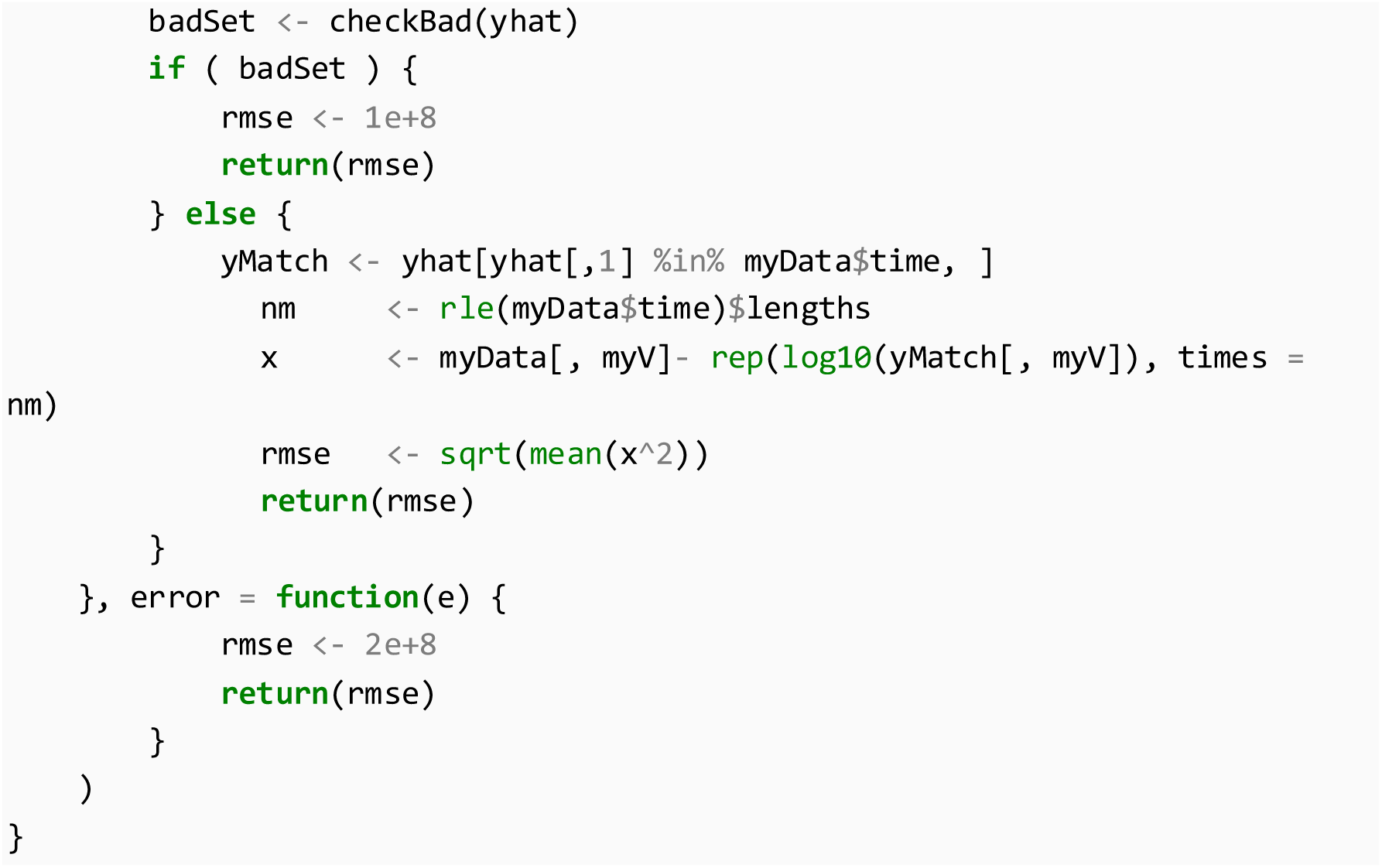
16. When a small sample size (*n*) is used or a large number of parameters are estimated (*k*), a corrected version of AIC should be used (23), i.e., AIC_C_ = AIC + 2*k*(*k* + 1) / (*n*–*k*+1).
17. Many parts of the codes can be speeded up by vectorizing and parallelizing the calculations, e.g., using *mcapply*, *snow* package. But different implementations are needed for Windows compared to Linux and Mac OS. Here, using base R codes the calculations can be applied to all operating systems.
18. Many innovative optimizing algorithms exist and continue to be improved. However, there is not any algorithm that can improve the estimation quality in the scarce and sparse of data (24). Therefore, it is recommended to seek for improvements in extra data sources instead of in variations of the other optimization techniques.
19. Using R-script editor is straightforward. However, a text editor tailored for programming will minimize coding errors and speed up considerably the process. Popular freeware editors for R include, but not limit to, R Studio, Sublime Text, Notepad++, Atom.
20. A complete R code as described in this chapter can be found at https://figshare.com/s/9f0c50984e470693839e

## Footnotes

§ Copy and paste the commands, press Enter (required internet connection).

